# Protein kinase gene declines linearly with elevation: a shared genomic feature across species and continents in lichenized fungi suggests role in climate adaptation

**DOI:** 10.1101/2022.11.02.514805

**Authors:** Dominik Merges, Francesco Dal Grande, Henrique Valim, Garima Singh, Imke Schmitt

## Abstract

Intraspecific genomic variability affects a species’ adaptive potential towards climatic conditions. Variation in gene content across populations and environments may point at genomic adaptations to specific environments. The lichen symbiosis, a stable association of fungal and photobiont partners, offers an excellent system to study environmentally driven gene content variation. Many species have remarkable environmental tolerances, and often form populations in different climate zones. Here we combine comparative and population genomics to assess the presence and absence of genes in high elevation and low elevation genomes of two lichenized fungi of the genus *Umbilicaria*. The two species have non-overlapping ranges, but occupy similar climatic niches in North America (*U. phaea*) and Europe (*U. pustulata*): high elevation populations are located in the cold temperate zone and low elevation populations in the Mediterranean zone. We assessed gene content variation along replicated elevation gradients in each of the two species, based on a total of 2050 individuals across 26 populations. Specifically, we assessed shared orthologs across species within the same climate zone, and tracked which genes increase or decrease in abundance within populations along elevation. In total, we found 16 orthogroups with shared orthologous genes in genomes at low elevation and 13 at high elevation. Coverage analysis revealed one ortholog that is exclusive to genomes at low elevation. Conserved domain search revealed domains common to the protein kinases (PKs) superfamily. We traced the discovered ortholog in populations along five replicated elevation gradients on both continents. The protein kinase gene linearly declined in abundance with increasing elevation, and was absent in the highest populations. We consider the parallel loss of an ortholog in two species and in two geographic settings a rare find, and a step forward in understanding the genomic underpinnings of climatic tolerances in lichenized fungi. In addition, the tracking of gene content variation provides a widely applicable framework for retrieving biogeographical determinants of gene presence/absence patterns. Our work provides insights into gene content variation of lichenized fungi in relation to climatic gradients, suggesting a new research direction with implications for understanding evolutionary trajectories of complex symbioses in relation to climatic change.

## Introduction

The intraspecific genomic variability substantially affects a species’ adaptive potential to ecological interactions and climatic conditions (Badet et al., 2020; Hirsch et al., 2014; Resl et al., 2022). Variation in genomic content at the gene-level, i.e. presence/absences of genes, within species are regularly found in bacteria, and are often associated with adaptations to specific environments and antibiotic resistances (Heuer et al., 2011; Masignani et al., 2005). Recent studies suggest that eukaryotic species, like bacterial ones, can have intra-specific variation in genomic content at the gene-level (Badet et al., 2020; Gerdol et al., 2020; Hirsch et al., 2014). To this date, it is largely unknown how relevant such variation in gene content is for eukaryotes, and if it is ecologically important for adaptations to specific environments. Intra-specific variation in gene content contributes to the formation of genetically diverse populations, and therefore its characterization is vital to understand the mechanisms of adaptation to specific environments (Drott et al., 2021; Plissonneau et al., 2018).

The lichen symbiosis, a stable association of mainly fungal and photobiont partners as well as an associated microbiome, lends itself to the study of environmentally driven gene content, because many species have remarkable environmental tolerances and maintain populations in different climate zones (Grimm et al., 2021; Jung et al., 2021; Kappen, 2000; Medeiros et al., 2021; Sierra et al., 2020; Singh et al., 2017; Tanunchai et al., 2022; Werth & Sork, 2014). Population genomic analyses based on single nucleotide polymorphisms (SNPs) suggest the presence of genome-wide differentiation between populations in different climate zones (Dal Grande et al., 2017; Rolshausen et al., 2022). Differentiation at the level of SNPs is often significantly correlated with differences in geography and ecology and may thus be involved in environmental specialization (Castillo et al., 2012; Hodkinson et al., 2012; Peksa & Skaloud, 2011). The only study which assesses variation in gene content associated with environmental adaptation is limited to a single species: Singh et al. (2021) show that some gene clusters associated with natural product biosynthesis in *Umbilicaria pustulata* have elevation-specific distributions. It is currently unknown 1) whether different species of lichen-forming fungi maintain homologous population-specific genes along environmental gradients and 2) whether intra-specific variation in gene content contributes to environmental tolerance of lichens.

Modern evolutionary approaches leverage DNA sequencing to infer ecological and evolutionary processes that occur at the population level. Here we combine comparative genomics and population genomics to assess the presence/absence of genes in high elevation and low elevation genomes of two lichenized fungi of the genus *Umbilicaria* (*Umbilicaria phaea* and *U. pustulata*). The two species have non-overlapping ranges but occupy similar climatic niches in North America (*U. phaea*) and Europe (*U. pustulata*), i.e. the cold temperate and the Mediterranean zones. We tracked gene content variation along replicated elevation gradients in both species based on a total of 2050 individuals in 26 populations. Specifically, we addressed the following research questions: a) Which genes are linked to environmental conditions at high elevation (cold temperate climate) and low elevation (Mediterranean climate) across species? To address this question, we assessed which genes are exclusive to climate zones. b) Do abundances of genes specific to a climate zone co-vary with elevation in populations of *U. phaea* and *U. pustulata*? To address the second question, we assessed which genes increase or decline in abundance with increasing elevation.

## Material and methods

### Study site and sample collection

We sampled 15 *U. pustulata* populations along three elevational gradients in Spain and Italy and 11 *U. phaea* populations along two elevational gradients in California, USA (Merges et al., 2021; Singh et al., 2022). Details of the sampling of European material are described in Singh at al. (2022) and the details of sampling of the North American material in (Dal Grande et al., 2017; Merges et al., 2021). Briefly, two of the European gradients are located in Central Spain, Sierra de Gredos, and one on the island of Sardinia. We collected fragments of 100 individuals each, at Mount Limbara (Sardinia, Italy; 6 populations, IT), Sierra de Gredos (Sistema Central, Spain; 6 populations, ESii) and Talavera-Puerto de Pico (Sistema Central, Spain; 3 populations, ESi), as described in Dal Grande et al. (2017). The Californian gradients are spatially separated by approx. 700 km. We collected fragments of 50 individuals each, at four populations along the Sierra Nevada gradient (38.084, −129 120.484) and at seven population along the Mt. Jacinto gradient (33.435, −116.484). We additionally collected four whole lichen thalli, one low-altitude individual from the Sierra Nevada population Uph16 and one from the high-altitude population Uph19, as well as a low-altitude and a high-altitude individual from populations ESii1and Esii6 of the Sierra des Gredos gradient for the reconstruction of reference genomes using PacBio Sequel II data.

### DNA extraction for population pooled sequencing

Genomic DNA was extracted separately from each fragment from all populations using a CTAB-based method (Cubero & Crespo, 2002; Dal Grande et al., 2017). Further, we created a pooled sample for each population containing equal amounts of DNA from each sample (i.e., Pool-seq; Dal Grande et al., 2017). Novogene Co., Ltd. (Cambridge, United Kingdom) and performed the library preparation (200–300 bp insert size). Libraries were sequenced on an Illumina HiSeq2000 with 150 bp paired-end chemistry at ∼90x coverage per population.

### DNA extraction for genomic sequencing

Genomic DNA for genome sequencing was extracted from dry thallus material of two samples of the same species (i.e. *U. phaea* or *U. pustulata*) collected in different climatic zones (i.e., low elevation/Mediterranean climate zone and high elevation/temperate climate zone). Lichen thalli were thoroughly washed with sterile water and checked under the stereomicroscope for the presence of possible contamination or other lichen thalli. DNA was extracted from all of the samples using a cetyltrimethylammonium bromide (CTAB)-based method (Mayjonade et al., 2016) as presented in Merges et al. (2021).

### PacBio library preparation and sequencing

SMRTbell libraries were constructed according to the manufacturer’s instructions of the SMRTbell Express Prep kit v2.0 following the Low DNA Input Protocol (Pacific Biosciences, Menlo Park, CA). Total input DNA was approximately 140 ng and 800 ng, respectively. Ligation with T-overhang SMRTbell adapters was performed at 20°C overnight. Following ligation, the SMRTbell library was purified with an AMPure PB bead clean up step with 0.45X volume of AMPure PB beads. Subsequently a size-selection step with AMPure PB Beads was performed to remove short SMRTbell templates < 3kb. For this purpose, the AMPure PB beads stock solution was diluted with elution buffer (40% volume/volume) and then added to the DNA sample with 2.2X volume. The size and concentration of the final libraries were assessed using the TapeStation (Agilent Technologies) and the Qubit Fluorometer with Qubit dsDNA HS reagents Assay kit (Thermo Fisher Scientific, Waltham, MA). Sequencing primer v4 and Sequel® II Polymerase 2.0 were annealed and bound, respectively, to each SMRTbell library. SMRT sequencing was performed on the Sequel System II with Sequel II Sequencing Kit 2.0 in “continuous long read” (i.e. CLR) mode, 30 hour movie time with no pre-extension and Software SMRTLINK 8.0. One SMRT Cell was run for each sample.

### De novo assembly of PacBio metagenomic sequence reads

We largely followed the pipeline described in Merges et al. (2021). In summary, we generated HiFi reads from the Pacbio Sequel II run using the PacBio tool CCS v5.0.0 with default parameters (https://ccs.how). Metagenomic sequence reads were assembled into contigs using the long-read based assembler metaFlye v2.7 (Kolmogorov et al., 2019). The assembled contigs were scaffolded with LRScaf v1.1.12 (https://github.com/shingocat/lrscaf; (Qin et al., 2019)). To retrieve the mycobiont genome, the received scaffolds were taxonomically binned via blastx using DIAMOND (--more-sensitive --frameshift 15 –range-culling) on a custom database (Singh et al., 2022) and the MEGAN6 Community Edition pipeline (Buchfink et al., 2014; Huson et al., 2007). The completeness of the genomes represented by the binned Ascomycota scaffolds was estimated using Benchmarking Universal Single-Copy Orthologs (BUSCO) analysis in BUSCO v4 (Simão et al., 2015) using the Ascomycota dataset.

### Pool-seq data processing

We filtered the pool-seq data for reads shorter than 80 bp, reads with N’s, and reads with average base quality scores less than 26 along with their pairs, and discarded them. We mapped the trimmed paired-end reads of each pool to the database of the identified genes using bowtie2 v2.4.1 (Langmead & Salzberg, 2012), using the flags: --very-sensitive-local, --no-mixed, --no-unal, --no-discordant.

### Gene prediction and genome annotation

Functional annotation of genomes, including genes and proteins (antiSMASH; antibiotics & SM Analysis Shell, v5.0) was performed with scripts implemented in the funannotate pipeline (Blin et al., 2017; Palmer & Stajich, 2019). First, the genomes were masked for repetitive elements, and then the gene prediction was performed using BUSCO2 to train Augustus and self-training GeneMark-ES (Borodovsky & Lomsadze, 2011; Simão et al., 2015). Functional annotation was done with InterProScan (Quevillon et al., 2005), egg-NOG-mapper (Huerta-Cepas et al. 2019, 2017) (Huerta-Cepas et al., 2017, 2019), and BUSCO v 5.1.2 (Simão et al., 2015) with ascomycota_db models. Secreted proteins were predicted using SignalP (Armenteros et al., 2019) as implemented in the funannotate “annotate” command. Proteins where further characterized by NCBI conserved domain search (Lu et al., 2020).

### Assessing gene content variation in the assembled fungi genomes

To identify presence/absences patterns of genes, we identified orthologs using orthoFinder (Emms & Kelly, 2015, 2019). OrthoFinder provides the most accurate ortholog inference method on the Quest for Orthologs benchmark test (Emms & Kelly, 2015, 2019). In orthoFinder (v.2.5.4), we assigned all genes to orthogroups using protein homology and constructed a pangenome of all four complete genomes (Badet et al., 2020). Shared orthologs (i.e. members of the some orthogroup) of low elevation (warm adapted) and high elevational (cold adapted) genomes were extracted using R v3.6.1 (R Core Team, 2019).

### Validating presence/absence of genes at population level

To validate population level gene presence or absence, we estimated the abundance of each ortholog in the low elevation (warm adapted) and high elevation (cold adapted) population based on the median coverage of pool-seq reads associated to each ortholog contig. Specifically, we used samtools (v1.15) depth to estimate the coverage of each basepair within the contig (Danecek et al., 2021). We assessed and visualized the data in R v3.6.1 (R Core Team, 2019).

### Gene distribution across Umbilicaria populations

Bowtie2 (v2.2.2) was used to map pool-seq reads to all ortholog contigs (using default settings). The number of mapped reads was counted per sample and normalized by dividing the number of mapped reads by the total read number of the respective sample to account for differences in sequencing depth. We modelled gene abundance (i.e., normalized read count) as a function of elevation using linear models. Linear models were fitted and plotted in R v3.6.1 (R Core Team, 2019).

## Results

### HiFi metagenomic sequencing reads of mycobiont

We reconstructed metagenomic sequences from a low-elevation and a high-elevation specimen of *U. pustulata* and *U. phaea*. Sequence output and quality for *U. pustulata* were summarized in Singh et al. 2022, for *U phaea* in Merges et al 2021 and in Table S1.

### Altitude-specific genes in the de-novo assembled genomes

We screened the de-novo assembled genomes of the low- and high-altitude samples for altitude-specific genes (Figure 1). Orthofinder revealed 16 orthogroups with shared orthologous genes (0.2 % of the total orthogroups) in low-elevation genomes and 13 in high-elevational genomes (0.1 % of the total orthogroups).

**Figure 1:**
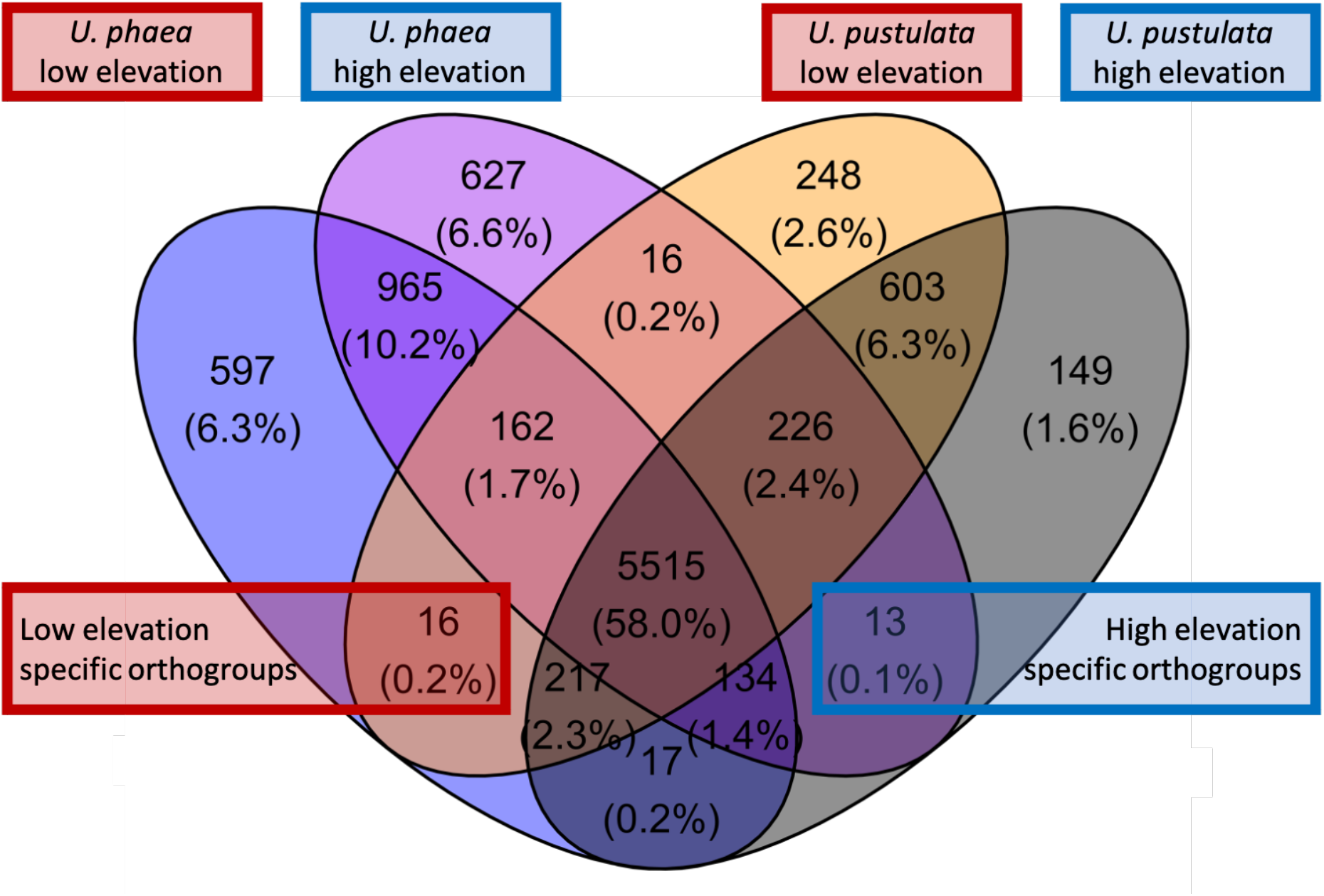
Venn diagram displaying orthogroups of *U. phaea* and *U. pustulata*. Red box highlights orthologs of the warm adapted (low elevation) *U. phaea* and *U. pustulata* genomes and the blue box of the cold adapted (high elevation) genomes.

### Presence/absence of genes at population level

To verify the presence/absence of detected orthologs, the coverage of each ortholog was calculated for the respective population at low and high elevation. The coverage analysis revealed one ortholog present in the genomes at low elevation to be consistently absent in populations at high elevation (Fig. 2.). The amino acid sequences of the ortholog could not be functionally annotated using the funannotate pipeline and was classified as “hypothetical protein”. NCBI’s conserved domain search revealed an alignment with the catalytic domain of Protein Kinases superfamily member PKc cd00180 (accession cl214531) as well as seven Tetratricopeptide repeats, indicating putative protein binding surfaces.

**Figure 2:**
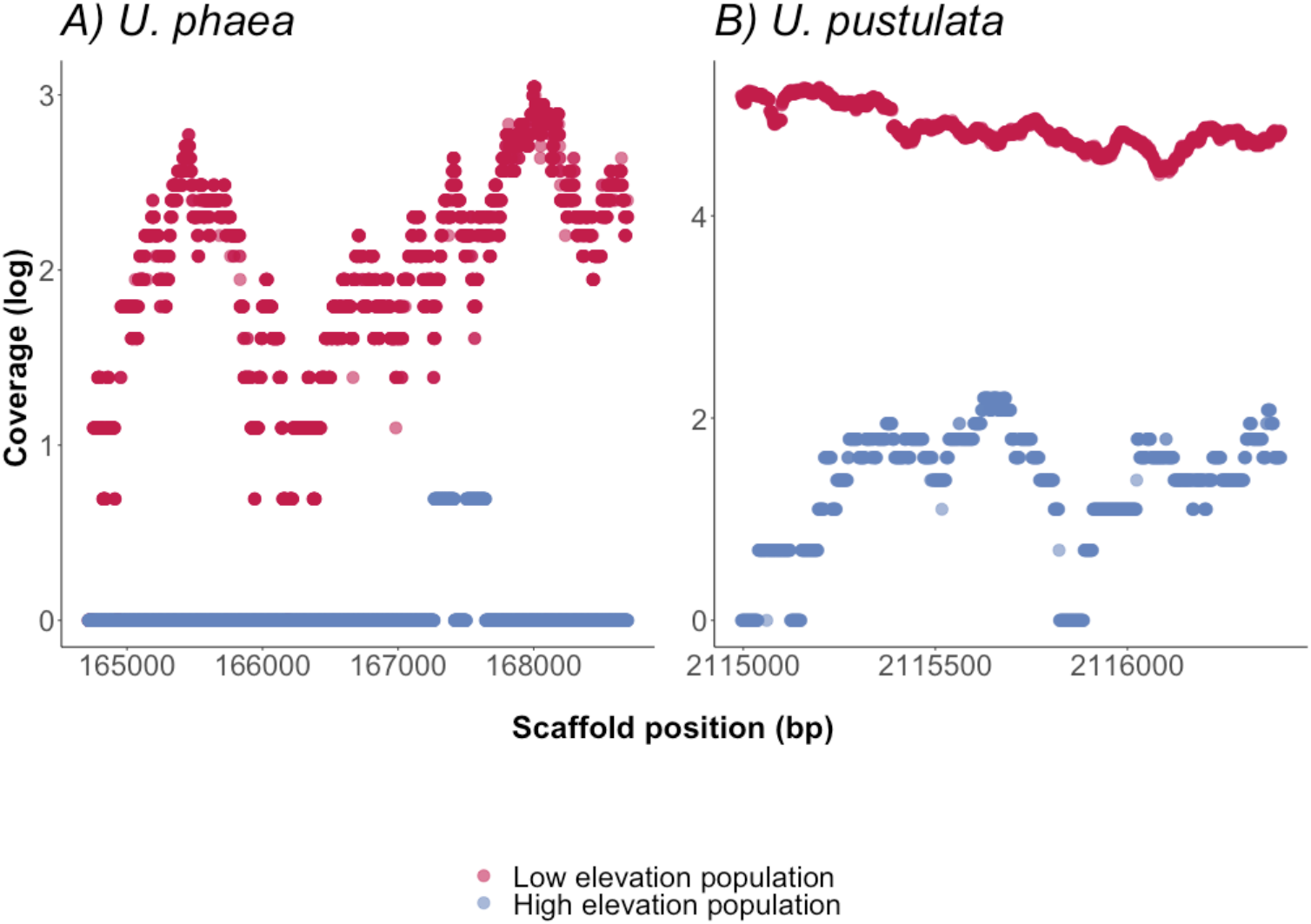
Presence/absences of orthologs in four de novo sequenced genomes of *U. phaea* and *U. pustulata* were verified by assessing the scaffold coverage in the warm adapted (low elevation) and the cold adapted population (high elevation) respectively. A) Coverage of ortholog in *U. phaea*: High coverage in warm adapted (low elevation) population and no coverage in high elevational population. B) Coverage of ortholog in *U. pustulata*: High coverage in in warm adapted (low elevation) population.

### Gene abundance distributions along gradients

The normalized read number of the identified orthologs, annotated as members of the Protein Kinases superfamily, showed a decline with increasing elevation across all populations (Figure 3).

**Figure 3:**
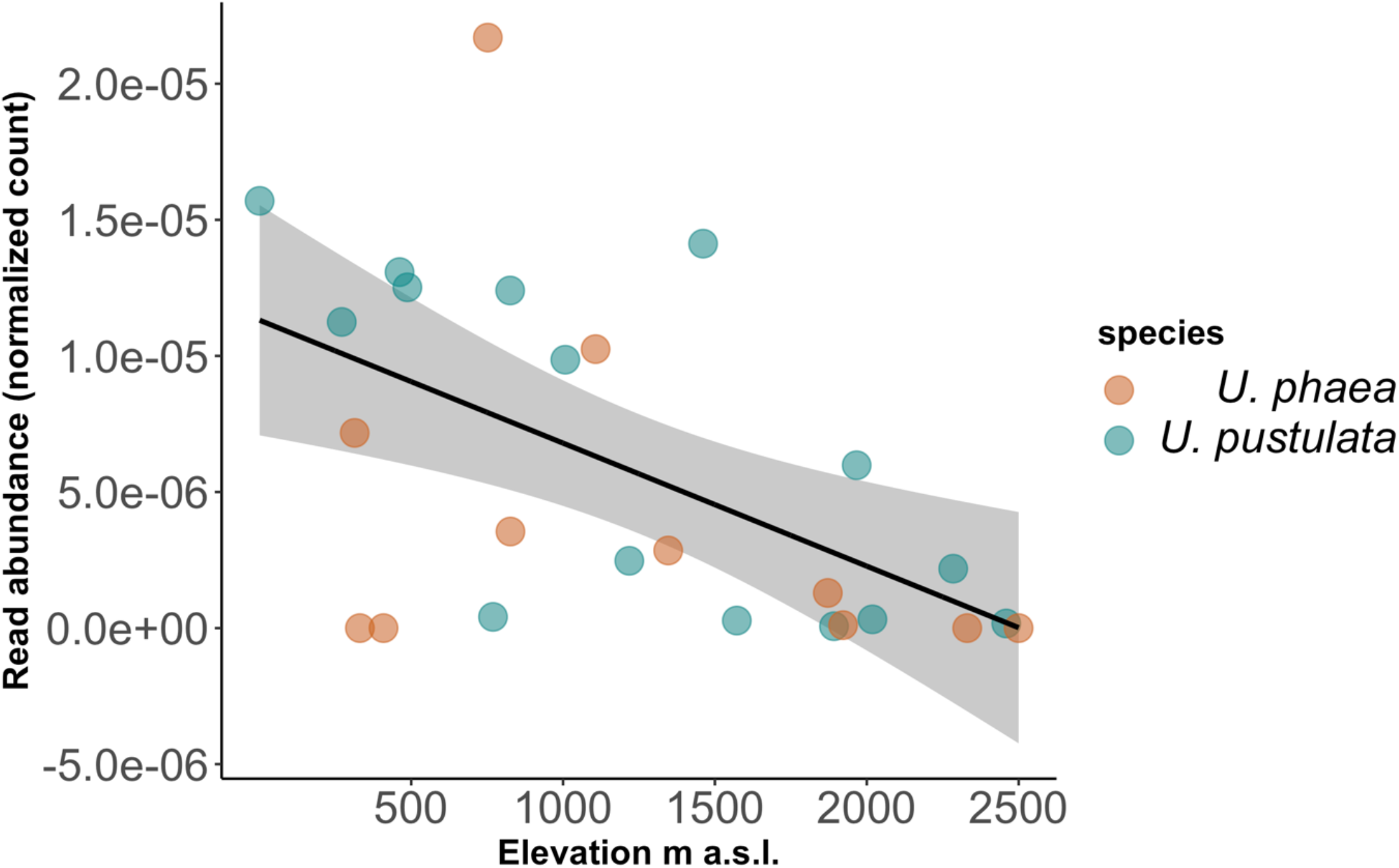
Read abundance of identified ortholog decreases significantly with in increasing elevation across *U. phaea* (brown circles) and *U. pustulata* (blue circles) populations.

## Discussion

Although adaptations to environmental gradients may lead to variation in gene content, assessments of gene presence/absence patterns across populations and species of lichenized fungi are still missing. In this study, we take an in-depth look at gene presence and absences in lichenized fungi of the genus *Umbilicaria*, and trace the discovered genes in lichen populations along five replicated elevation gradients across two continents. While our whole genome comparison based on four de novo sequenced specimen (two per species) suggested up to 29 elevation-specific orthogroups, the population-level verification approach showed only one gene, putatively encoding a protein kinase (PKs), which linearly declined in abundance with increasing elevation, and was truly absent in the highest population. This suggests either high strain-specificity of certain genes, or high false positive recovery of gene presence/absence patterns when relying on comparative genomics approaches based on only a few individuals. Such approaches can be potentially misleading when interpreting the evolutionary significance of gene content variation at population level. Regarding the PKs gene consistently absent in high elevation genomes and populations, we found that the discovered gene declines linearly across all populations, suggesting an evolutionary benefit only at lower altitudes. Alternatively, the loss of the gene at higher elevations might benefit individuals in cold climates. To our knowledge, we report for the first-time parallel gene presences and absences patterns correlating with climatic niches in different species of lichenized fungi. However, it remains unclear if this is an adaptive trait.

In bacteria variation in gene content is assumed to be driven by selection for environmental conditions that are relatively rare across the entire range of a species (Qi et al., 2017). Recent evidence suggests that specific populations of lichenized fungi may contain unique biosynthetic gene clusters (Singh et al. 2021), and our current findings show that also other genes can be elevation-specific. The gradual gene loss across populations with increasing elevation may suggests a decline of selective benefit and may indicate that certain variations in gene content could be of functional importance for local adaptation and climatic tolerances in lichenized fungi. The conserved domain search revealed a catalytic domain of a PK, a common eukaryotic protein superfamily. PKs selectively modify other proteins by phosphorylation, changing their enzymatic activity, cellular location and association with other proteins (Asano et al., 2005; Cheng et al., 2002; Hanks, 2003; Heinisch & Rodicio, 2018). Within a genome, PKs are encoded by a large multigene family with genes being distributed among multiple chromosomes. Putatively, the high number of PK genes has arisen by genome segmental duplication events (Asano et al., 2005; Heinisch & Rodicio, 2018). In our study, the presence/absence of a single PK gene may suggest a climate-specific ancestral genome segmental duplication event. However, due to the scarcity of functional annotation of non-model organisms and the resulting lack of in-depth functional annotation of the gene in question, the mechanisms generating such population level diversification are yet to be understood.

While environmental adaptations are commonly highly polygenic (Barghi et al., 2019; Hartke et al., 2021; Pfenninger et al., 2021; Rivas et al., 2018), there is increasing evidence of the effect of single gene content variation (Liu et al., 2021). For example, as has been recently shown in agave, where a single gene encoding a phosphoenolpyruvate carboxylase enhances the plant’s climate resilience (Liu et al., 2021). Not only the gain of genes, but also the loss of genes has been associated with adaptive traits, such as the evolution of particular diets in bats (Blumer et al., 2022). Therefore, we consider the parallel loss of a homologous gene in two species and two geographic settings a rare find, and a step forward in understanding the genomic underpinnings of climatic tolerances in lichenized fungi. Future research should address the functional importance of the gene present at low altitude in the Mediterranean climate zone, and specifically explore the effects of variation in gene abundances across populations. Additionally, future research should consider using heterologous expression approaches to reveal whether the gene presence could induce tolerances to warm conditions.

## Conclusion

Our study demonstrates how comparative genomics in combination with population genomic data can detect patterns of gene content variation across climatic gradients. In addition, the tracking of gene content variation across populations provides a widely applicable framework for retrieving meaningful biogeographical determinants of gene presence/absence patterns. To this end our work provides insights into gene content variation of lichenized fungi in relation to climatic gradients. This suggests a promising new research direction with implications for understanding evolutionary trajectories in relation to climatic change.

## Data Accessibility

Raw sequence reads were deposited in the Sequence Read Archive under the BioProject PRJNA693984 and PRJNA820300.

## Acknowledgments

We thank Carola Greve and Jürgen Otte for the laboratory procedures, and Christoph Sinai, Tilman Schell and Anjuli Calchera (Frankfurt am Main) for support with bioinformatics. We thank Barbara Feldmeyer and Markus Pfenninger (Frankfurt am Main) for their valuable suggestions on analyses.

## Funding information

This research has been partly funded by the Hesse’s “state initiative for the development of scientific and economic excellence” (LOEWE initiative).

## Authors’ contributions

D.M., and I.S. conceived the ideas; I.S. and F.D.G. collected the data, D.M., G.S., H.M. performed genome assembly and annotations, D.M. analyzed data, F.D.G. provided analytical guidance; D.M. and I.S. wrote the manuscript. All authors contributed to the various drafts and gave final approval for publication.

## Supplement

**TABLE S1.**
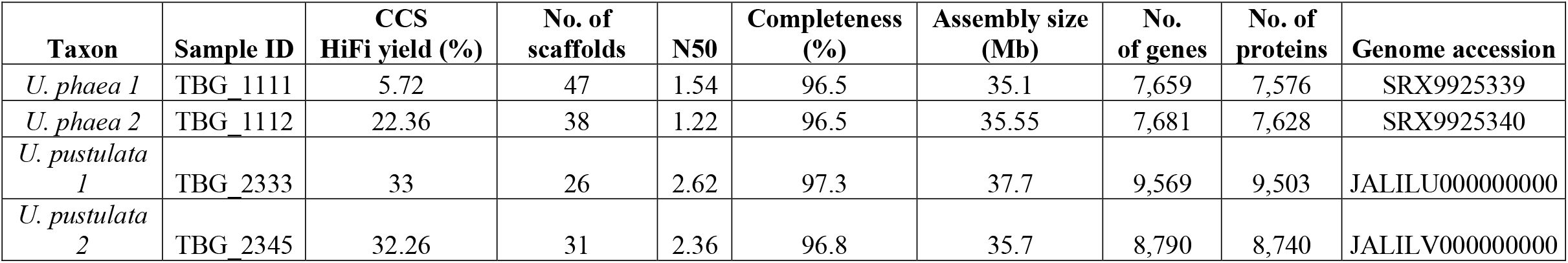
Genome quality and annotation statistics

## Notes

### Competing Interest Statement

The authors have declared no competing interest.

